# Multisensory integration of stimulus-driven and goal-driven signals during urgent saccadic choices

**DOI:** 10.64898/2026.06.18.733213

**Authors:** Ashley N Paro, Bashirul I Sheikh, Terrence R Stanford, Emilio Salinas

**Affiliations:** Department of Translational Neuroscience, Wake Forest University School of Medicine, 1 Medical Center Blvd., Winston-Salem, NC 27157-1010, USA

**Keywords:** antisaccade, attention, audition, crossmodal capture, decision making, multimodal, salience

## Abstract

The ability to orient or attend to sensory events is generally greater in response to visual and auditory cues occurring together than in response to single-modality cues occurring alone. In such cases the perceptual fusion of cross-modal stimuli (multisensory integration) depends on low-level features (e.g., location, intensity) and follows well established principles. However, less is known about multisensory integration mechanisms when behavioral responses are less direct and require top-down control. Here we investigate this in human participants using an urgent multisensory choice task that effectively dissociates stimulus-driven and goal-driven contributions to performance based on their distinct temporal signatures. Task conditions varied the modality of the cues (auditory, visual, or both), their location (left or right), and the rule defining the correct choice (look toward or away from a given cue). When spatially coincident cues were associated with the same response rule (“look away”), we observed multisensory enhancement and performance remained close to a statistical expectation as the choice process unfolded. However, when spatially disparate cues were associated with different rules but the same target, one cue dominated performance and the other produced crossmodal capture, i.e., low-level competition. The results indicate that the efficacy of multisensory integration is dictated by the stimulus-and goal-driven signals produced by each cue, with all four factors rapidly interacting in accordance to the dynamics of spatial attention.

**Significance Statement:** Auditory and visual stimuli located near each other in space and time are typically bound into a single sensory percept that draws attention most effectively. However, it is unclear whether such “multisensory integration” occurs during behaviors that go beyond directly attending or orienting to cue stimuli and require top-down control. We investigated this using a novel task design with which stimulus-driven and goal-driven contributions to performance can be accurately identified. We found that multisensory enhancement depends not so much on the complexity of the requested cue-response associations, but rather on the timing and alignment of the stimulus-and goal-driven signals derived from each cue (auditory and visual) — similar to the way that such signals dictate the allocation of spatial attention.

## Introduction

Natural events in the environment often yield sensory signals from multiple modalities. As we see a door moving ever so slightly and hear a faint metal creak, we may suspect that the cat has sneaked into the room. Together, the movement and the sound generate a stronger, more reliable signal than either stimulus alone. Many studies in so-called “multisensory integration” have aimed to characterize and quantify the resulting behavioral benefit (Miller, 1986; Corneil et al., 2002; Burnett et al., 2004; Diederich and Colonius, 2004; Drugowitsch et al., 2014) or the underlying neurophysiological substrates (Stanford et al., 2005; Stein and Stanford, 2008; Angelaki et al., 2009) whereby inputs from different modalities are merged into a more effective multisensory output. This has resulted in a conceptual framework that accounts for performance based on specific neural operations (Ma and Pouget, 2008). This framework includes a number of heuristic principles that encapsulate the conditions under which multisensory stimuli may yield a performance benefit (Meredith et al., 1987; Meredith and Stein, 1996; Stein et al., 1989, 2009), such as temporal proximity, spatial congruence, and inverse effectiveness (the benefit is larger when the unisensory signals are similarly weak).

However, all this is generally applicable to situations involving relatively simple behaviors, such as overtly orienting toward a salient stimulus or covertly attending to it (without shifting gaze). Behavioral responses in such cases are strongly stimulus-driven, i.e., fast and reflexive; indeed, key underlying neural integration mechanisms are observed under anesthesia (Wallace et al., 1989). Less is known about multisensory integration when behavioral responses require some type of endogenous control. This case is more complex not only because endogenous control can take many forms, but also because stimulus-driven contributions are never absent, and their in-fluences on ostensibly goaldriven behavior are difficult to isolate. A critical point in this regard is that the conditions for producing multisensory enhancement may not be the same for exogenous and endogenous signals, and the degree to which such signals are sympathetic or antagonistic could have a profound influence on the resulting behavior. With this fact in mind, here we use a new set of urgent-choice tasks to determine how the interaction between stimulus-driven and goal-driven factors dictates behavioral outcomes when goal-driven choices are based on multisensory cues.

Imposing urgency turns out to be extremely useful for dissociating exogenous and endogenous contributions to visuomotor behavior (Salinas et al., 2019; Poth, 2021; Goldstein et al., 2022, 2024; Oor et al., 2023, 2025; Zhu et al., 2024; Kattner et al., 2026; Krause and Poth, 2025). Choices made under time pressure may evolve to various endpoints, ranging from entirely uninformed guesses (near chance) to fully informed decisions (approaching 100% correct), and the transition between them provides a direct readout of the timecourse of the underlying perceptual or cognitive evaluation (Stanford et al., 2010; Stanford and Salinas, 2021). In general, this approach clearly identifies choices that are exogenously guided (early, involuntary, stimulus-dependent) from those that are endogenously guided (late, voluntary, goal-driven), which is a fundamental functional distinction (Corbetta and Shulman, 2002; Theeuwes, 2010; Carrasco, 2011; Noyce et al., 2023).

Here we investigate saccadic choices that are (1) urgent and (2) based on multisensory (visual and auditory) stimuli. The confluence of these two properties is illuminating both from the perspective of the mechanisms whereby stimulus information from different modalities can be combined to yield a stronger, more reliable target representation, and from the perspective of the attention mechanisms whereby different potential targets are prioritized. Two sides of the same behavioral coin (Driver and Spence, 1998; Koelewijn et al., 2010; Santangelo and Macaluso, 2012). We find the effect of combined multisensory cues on a rule-driven choice to be highly context-dependent, yet readily understood as the temporally weighted superposition of two processes: an early, stimulus-driven one reflecting the spatial congruence of the unisensory cues, and a later, goal-directed one reflecting the dynamics of endogenous attention.

## Materials and Methods

### Subjects and setup

Experimental participants were 15 healthy human volunteers 23–62 years of age (median age was 37); all but one were female. They were recruited from the Atrium Health Wake Forest Baptist Medical Center and broader Winston-Salem communities. All participants had normal or corrected-to-normal vision and normal hearing. All provided written informed consent prior to the experiment and were compensated for their participation. Of the 15 participants, 12 completed all 3 task variants, two (numbers 2 and 10) completed only two of the variants, and one (number 3) abandoned the study without completing any of the variants. We only consider the data of the remaining 14 participants. All experimental procedures were approved by the Institutional Review Board of Wake Forest University School of Medicine in accordance with protocol IRB00097510.

Similar to previous studies (Goldstein et al., 2022, 2024), all experimental sessions were conducted in a private unlit room. Participants were seated on an adjustable chair with their forehead and chin supported. Visual stimuli were presented on a 24 inch VIEWPixx LED monitor (VPixx Technologies Inc., Saint Bruno, Quebec, Canada; 1920 x 1200 screen resolution, 120 Hz, refresh rate, 12-bit color) placed approximately 57 cm from the participant. Auditory stimuli were emitted from custom-made speakers positioned to the left and right of the screen and horizontally aligned with the visual stimuli. Sounds were triggered by the VIEWPixx monitor and amplified using a 2×75 Watt PYLE Pca3 Stereo Power Amplifier. Eye position was monitored monocularly and recorded using an EyeLink 1000 infrared camera and tracker (SR Research, Ottawa, Canada; 1000 Hz sampling rate). All stimulus presentation and data collection routines were implemented using Matlab (Mathworks, Natick, MA) and the Psychtoolbox 3.0 package (Brainard, 1997; Kleiner, 2007) running on a standard Windows-based desktop computer.

### Basic task design

We ran three variants of the compelled multisensory choice (**CMC**) task. These differed depending on the task goal as well as on the modality and spatial congruence of the cue stimuli presented. For all task variants and trial types the sequence of events was as follows.

First, a stimulus appeared in the middle of the display and the participant fixated on it. The color and shape of the fixation stimulus served as a reminder of the task variant and associated instructions (see below), which remained fixed for each block of (100) trials. After 300 ms of fixation two open targets appeared, one to the left and one to the right of the central fixation stimulus (Targets on). These targets indicated two things: the locations where a visual cue could appear (if such a cue was to be presented in the trial) and the desired endpoints of the choice saccades. After a short delay (300, 400, or 500 ms) the central fixation stimulus disappeared (Go), instructing the participant to make an eye movement. Although at this point the correct choice was still unknown, the participant had to attempt to respond right away in order to meet a reaction time (**RT**) deadline (350–450 ms adjusted for each participant). The sensory cue indicating the correct choice was revealed after the go signal (Cue), once a period of time called the gap had elapsed. The gap varied randomly between trials, typically ranging from 0 ms (cue simultaneous with go signal) to 250 ms (typical saccadic RT). Negative or “easy” gaps (−300 or −200 ms) were also used with some participants. Negative gaps correspond to trials in which the cue stimulus is presented before the go signal, so more time is available for processing the cue. Visual and or auditory cues remained until the participant responded by making an eye movement to one of the open targets (Saccade).

To make sure that the participants responded on time, feedback was provided after each trial; such feedback was never indicative of the correct saccade direction. Some participants (participants 1–11) were given negative feedback; for them the words “Go faster!” appeared on the center of the screen whenever the RT limit was exceeded. The rest of the participants (participants 12–15) were given positive feedback (the open targets turned from gray to green whenever the RT limit was met). Both approaches helped the participants adjust their response urgency. The task was run in blocks of 100 valid trials, i.e., trials that were completed within the required RT window.

The fixation stimuli were small (0.5° diameter), low-luminance, centrally located shapes that differed depending on the task variant (see below). Saccade choice targets were large (2.5 diameter), gray, outlined circles placed along the horizontal line 8° to the left and right of the central fixation point. Visual cue stimuli were small (0.5° diameter) gray spots with blurred edges. Each presented visual cue was displayed inside one of the peripheral choice targets and remained there until a saccade was made. Auditory cue stimuli were white noise bursts that lasted until a saccade was made.

Visual and auditory modalities transduce different forms of energy, and for many years a “prepotency of visual information” has been recognized (Colavita, 1974; Spence et al., 2012; Li et al., 2017; Malevich et al., 2026). With this in mind, to balance the likelihood that either sensory modality could be picked as the target of a saccade, the luminance of the visual cues was set close to detection threshold (0.22 cd/m^2^ for 11 participants, with RGB vector 0.025× [1 1 1]; 2.8 cd/m^2^ for 3 participants, with RGB vector 0.15×[1 1 1]) whereas the intensity of the auditory cues was kept well above threshold (the amplifier output was set to 38% of its maximum, which produced a noise burst of ∼75 db).

### Task variants

Trials could be either unisensory (only one cue stimulus was presented, visual or auditory) or multisensory (both visual and auditory cues were presented). For all task variants unisensory and multisensory trials were interleaved within a block. The trial types and cue locations were random, so participants could not predict the correct choice or whether one or two stimuli would be presented. Multisensory trials could be either spatially congruent (cue stimuli presented on the same side) or spatially incongruent (cue stimuli presented on opposite sides); however, congruent and incongruent trials were never interleaved within a block. The three task variants or experimental blocks were as follows.

During blocks of congruent antisaccades (**CA**) the fixation stimulus was a pink triangle and the rule was “look away from any visual or auditory cue.” Three trial types were interleaved: unisensory visual (V), unisensory auditory (A), and congruent multisensory (V+A), in which a visual and an auditory cue originated from the same side. The correct response was a saccade to the open target diametrically opposite to whichever cue stimulus was presented.

During blocks of incongruent visual antisaccades (**IVA**) the fixation stimulus was a pink circle and the rule was “look away from the visual cue.” Two trial types were interleaved: unisensory visual (V) and incongruent multisensory (V(A)), in which a visual and an auditory cue originated from opposite sides. The correct response was a saccade to the open target diametrically opposite to the visual cue, which was always presented.

During blocks of incongruent auditory prosaccades (**IAP**) the fixation stimulus was a green square and the rule was “look toward the auditory cue.” Two trial types were interleaved: unisensory auditory (A) and incongruent multisensory (A(V)), in which an auditory and a visual cue originated from opposite sides. The correct response was a saccade to the open target on the same side as the auditory cue, which was always presented.

During their first session, participants completed several practice blocks (of 20 trials) to get acquainted with the stimuli and timing requirements of the task. Thereafter such practice runs only occurred whenever a new variant was introduced. All participants performed the three variants in the same sequence: IVA first, then IAP, and CA last.

## Data processing

A participant made a saccadic choice when their gaze left a central window (2^◦^ radius) around the fixation stimulus. The exact direction of the choice saccade was calculated offline together with its onset time (using a velocity criterion of 4 /s). For each choice the RT was computed as the interval between the go signal and the onset of the corresponding response saccade. Saccades were excluded from analysis if they occurred before the go signal (fixation breaks), if they were predominantly vertical, if the RT was too long (*>* 800 ms), if a blink occurred within 100 ms of saccade onset, or if the eyes drifted outside of the fixation window without exceeding the velocity threshold. The vast majority of excluded trials (∼85%) were fixation breaks; others accounted for approximately 3% of the collected data.

Over the course of 6 or 7 experimental sessions, each participant completed 14–30 blocks of trials for each of the three task variants. Across the final 14 participants this resulted in a total of 85275 trials that were considered acceptable according to the criteria just described. The median number of acceptable trials per participant was 6149 (range, 4472–7469). For data analysis purposes what matters is the number of trials collected for each of the distinct trial types (7 in total across the 3 task variants), which yield a separate performance curve for each participant. As such, the collected data included approximately 878 trials per individual curve, which is sufficient for resolving timing differences of ∼10 ms between conditions at the individual participant level (Salinas et al., 2019; Goldstein et al., 2022, 2024).

Data were excluded in three instances in which participants were essentially unable to perform the task as required. The data from participant 1 were excluded from analysis of the CA experiment because they yielded performance in A trials that was close to chance at all PTs. The data from participants 1 and 10 were excluded from analysis of the IAP and IVA experiments, respectively, because they yielded extreme visual capture with virtually no recovery. In all cases, the performance of these participants was otherwise consistent with the rest of our sample, and inclusion or exclusion of these outlier data did not alter any results qualitatively.

## Data analysis

Data analyses were performed using Matlab (The MathWorks, Natick, MA) and Python. As in other urgent tasks (Poth 2021; Stanford and Salinas, 2021; Kattner et al., 2026; Krause and Poth, 2025), performance in the CMC task is fundamentally dictated by the processing time (**PT**), which is the maximum amount of time during which the participant can see or hear the cue in each trial before initiating a response. Accordingly, the critical performance metric in the CMC task is the tachometric curve, which describes the probability of making a correct choice as a function of PT. Tachometric curves were produced by determining the PT for each trial, sorting trials into PT bins (which shifted every 1 ms), and calculating the fraction of correct responses for all the trials inside each bin. Tachometric curves and related analyses were carried out both for individual participants and for their pooled data. Pooling means that all the choices from a group of participants were combined into one large dataset, as if produced by a single, aggregate participant. Tachometric curves for individual participants had a bin width of 80 ms; those for pooled data had a bin width of 40 ms.

## EXPERIMENTAL DESIGN

The basic idea for interpreting the data is to create separate tachometric curves for each of the 7 distinct trial types (found across all task variants) and then perform three types of comparison. Within each task variant, comparisons between unisensory and multisensory tachometric curves should indicate whether performance is facilitated or hindered by the presence of a second cue. Second, the effects may be different for short-PT trials, which are predominantly stimulus-driven, compared to long-PT trials, which are predominantly goal-driven. Finally, for any such comparisons, the results may also depend on congruence; in the short-PT regime, stimulus congruence (i.e., spatial alignment between cues) is likely to be most relevant, whereas in the long-PT regime, what matters the most may be either target congruence (i.e., spatial alignment between the saccade endpoints associated with each cue) or rule congruence (i.e., whether the two cues follow the same or different stimulus-response rules, which are “saccade to the stimulus” or “saccade away from the stimulus”). In combination, the 7 experimental conditions sampled should reveal key constraints or principles for how signals from different modalities are combined during goal-driven multisensory choices.

## PROBABILITY OF CAPTURE

The probability of capture (Kattner et al., 2026), or *P_CAP_*, was defined as

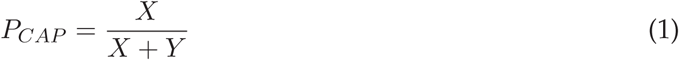

where *X* represents the area between the tachometric curve and the chance line when the curve is below said line, and *Y* is the area between the curve and the chance line when the curve is above. Defined this way, *P_CAP_* represents the proportion of erroneous choices that cannot be accounted for by chance; it measures the fraction of saccades that were directed toward the worse alternative above and beyond the fraction expected by chance. Thus, *P_CAP_* values of 0 and 1 correspond, respectively, to performance curves that either stay at or above chance, or stay at or below chance for the full PT range considered. Intermediate values result when the tachometric curve swings below and above chance. For each tachometric curve, *P_CAP_* was computed for PT values in the range 50–300 ms.

As done previously (Kattner et al., 2026), a confidence interval (**CI**) was calculated for each *P_CAP_* value by bootstrapping (Davison and Hinkley, 2006; Hesterberg, 2014). That is, for a given tachometric curve the underlying trial-wise data were resampled with replacement, *P_CAP_* was re-computed from the the corresponding (resampled) tachometric curve and saved, and the process was repeated many times (10000 iterations) to generate a distribution of *P_CAP_* values. Then, a 95% CI was obtained by calculating the 2.5 and 97.5 percentiles derived from the bootstrapped distribution. Note that *P_CAP_* is always positive or zero, by definition. Thus, from the same boot-strapped distribution, we also obtained the fraction of iterations for which *P_CAP_* was equal to zero. We report the corresponding probability, *P* (0), as an additional indicator of the significance of the measurement. It has a precision of 0.0001 given the number of iterations used.

## CURVE RISE POINT

For any tachometric curve the rise point was defined as the PT at which performance first exceeded a threshold value of 70% correct. Defined this way, the rise point indicates how quickly the cue stimulus is perceptually assessed. If one tachometric curve is simply shifted along the x-axis relative to another, the difference in rise points is equal to the relative shift between curves (in milliseconds of PT).

The method for calculating the rise point consisted of two steps. First find all the PTs for which the tachometric curve is near 70% correct (say, between 65% and 75%). Then, from those PTs, select the largest, continuous PT interval for which the curve values bordering the left edge of the interval are below the 65% limit and the values bordering the right edge are above the 75% limit. The rise point is then set equal to the midpoint of the selected PT interval. For curves that are relatively smooth, this procedure produces results that are nearly identical to those obtained by curve fitting (Shankar et al., 2011; Kattner et al., 2026), but it also works reasonably well when the data are noisy. For each rise point, a CI was calculated by bootstrapping in the same way as described above for *P_CAP_*.

## ASYMPTOTIC FRACTION CORRECT

Performance in the task was also quantified by calculating the mean fraction of correct choices at relatively long PTs for which the tachometric curve was consistently above chance. For this we first calculated the rise point of the tachometric curve and then identified all the trials with PTs higher than the rise point. The asymptotic fraction correct is simply the fraction of correct choices for those long-PT trials. This quantity is indicative of the reliability of performance for choices that were in all likelihood informed by the sensory cue(s). For some analyses we also computed the mean fraction correct for all the choices falling within a specific PT window (indicated in each case). A 95% CI was calculated for each fraction correct using binomial statistics; specifically the Agresti-Coull method (Agresti and Coull, 1998).

## PREDICTED MULTISENSORY PERFORMANCE

To quantify the effectiveness of cue integration, a novel statistical criterion was used to predict the performance on multisensory trials based on that observed during unisensory trials. The method is described in detail elsewhere (Salinas and Stanford, 2024; see also Coen et al., 2023; Oor et al., 2025), but the following is a brief overview.

The approach is based on the concept of conditional independence (Dawid, 1979). Two events *A* and *V* are conditionally independent relative to a third event *C* if their joint probability given *C* is such that

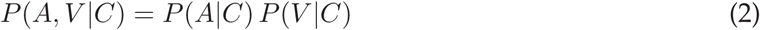

where *P* (*A, V* |*C*) is the probability that events *A* and *V* occur given that event *C* has occurred. In other words, if *C* is known, *A* and *V* occur independently of each other. An equivalent formulation of this concept is to say that, once condition *C* is known, the probability that *V* occurs is fixed regardless of whether *A* occurs or not. That is,

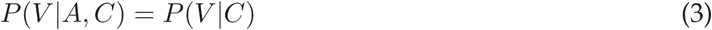

is an alternate, equivalent definition. Note that these expressions are distinct from the standard notion of independence, which for events *A* and *V* would correspond to

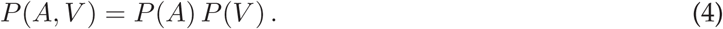

This condition is different and irrelevant to the problem at hand.

In the present study the event *C* corresponds to the outcome of a choice, which may be correct (*C* = 1) or incorrect (*C* = 0); event *A* corresponds to the presence (*A* = 1) or absence (*A* = 0) of an auditory cue; and *V* is the same but for a visual cue. Conditional independence is essentially a constraint on the three-way relationships between the three variables of interest as captured by their joint probability, *P* (*A, V, C*). Here, such constraint is useful because it means that the effect that *A* and *V* may have on *C* jointly (together) is fixed once their separate (individual) effects on *C* are known.

Specifically, the quantity of interest is the probability *P* (*C*|*A, V*), which describes how the likelihood of a correct outcome varies depending on which cues are shown. When *A* and *V* are conditionally independent with respect to *C*, Equation 2 can be used in combination with Bayes theorem to derive the following result

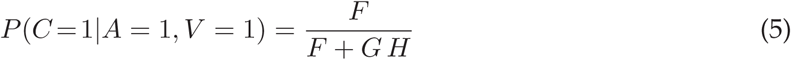

where we have defined the abbreviations

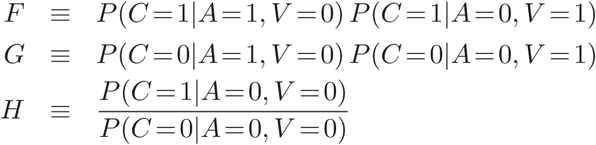

(Salinas and Stanford, 2024). Equation 5 is a prediction for the probability that the choice is correct given that both cues are presented. It is based on three quantities: the effect of *A* alone on choice outcome (represented by the terms conditioned on *A* = 1 and *V* = 0), the effect of *V* alone on choice outcome (represented by the terms conditioned on *A* = 0 and *V* = 1), and the probability of a correct choice when neither cue is shown (represented by the terms conditioned on *A* = 0 and *V* = 0). It is a parameter-free benchmark for quantifying how much knowledge is gained when the two cues are presented together rather than separately — if they do not interact. This prediction is to be contrasted with the empirical result for the probability of a correct choice measured when both cues are *actually* shown simultaneously.

For example, suppose that the probability of a correct choice given that only the auditory cue was presented is

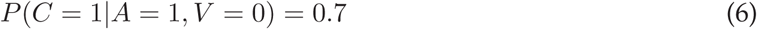

and that the probability of a correct choice given that only the visual cue was shown is

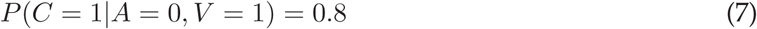

and also note that the probability of a correct choice given that no cue was shown is

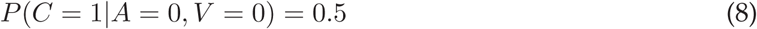

which corresponds to chance performance. Using these numbers, and recalling that *P* (*C* = 0|*A, V*) = 1 − *P* (*C* = 1|*A, V*), we can compute the terms *F*, *G*, *H* and use Equation 5 to find that *P* (*C* = 1|*A* = 1*, V* = 1) = 0.9; this is the prediction. It is larger than either of the probabilities conditioned on a single cue (Equations 6, 7) because they are in agreement (both above the chance level) and because together the two cues are statistically more informative of choice outcome. Comparisons with the experimental data obtained from actual multisensory trials are interpretable. Performance at or above the prediction would suggest that observed outcomes were guided by integrated signals, whereas performance below the prediction would suggest that observed outcomes were selected from separate potential outcomes.

To generate a predicted tachometric curve for multisensory trials in the CA variant, Equation 5 was applied at each PT bin. The required *F* and *G* terms were computed, also for each bin, using the tachometric curves from the unisensory auditory and visual trials. The *H* term was always equal to 1 because chance performance in the CMC task is constant at 50% correct. For the predicted probability at each bin, a distribution of likely values was generated using binomial statistics and the known numbers of A and V trials that went into the unisensory calculations at that bin. The resulting distribution for the predicted probability correct was used to derive 95% CIs.

## STATISTICS

For any performance metric (rise point, *P_CAP_*, fraction correct, etc.), differences between experimental conditions were assessed based on the 95% CIs obtained for each condition or on the underlying distributions (generated either by bootstrapping or via binomial statistics, as described above). For any such metric, we refer to mean values *m*_1_ and *m*_2_ as being “decidedly different” if there was no overlap between their 95% CIs. This overlap criterion provides a simple statistical shortcut for rapidly assessing differences betweeen multiple measurements. Numerical experiments indicate that there is a consistent correspondence between this rule and the traditional p-value for the difference between *m*_1_ and *m*_2_. That is, for adjacent 95% CIs with zero overlap, p-values are between 0.01 and 0.001 under a variety of sample sizes and distribution shapes, and p-values decrease further as the gap between CIs increases.

For the CA variant of the CMC task, we compared the predicted probability correct discussed in the previous subsection (*C_P_*) with that measured empirically (*C_E_*). For this, a significance p-value for the difference was calculated as follows (suppose *C_E_* − *C_P_ >* 0). First, distributions for the predicted and empirical values were obtained by bootstrapping, i.e., by resampling with replacement the trials underlying each curve, as described above. Pairs of samples (*c_E_*, *c_P_*) were then drawn repeatedly from these distributions, and the fraction of draws in which the result was opposite to that observed (*c_E_* ≤ *c_P_*) was taken as the significance of the observed difference. Because this number was based on 10000 iterations, or draws, 0.0001 is the minimum p-value reported.

## DATA SORTING FOR HISTORY ANALYSIS

History-conditioned tachometric curves were generated by considering choices that were preceded by cues of a specific modality. For instance, in the CA experiment the V trials were split into three groups: those that were preceded by a V trial, those that were preceded by an A trial, and those that were preceded by a V+A trial. A history-conditioned tachometric curve was generated for each of the three trial subsets. Similar sorting was applied to the data from other experiments. Comparison of history-conditioned curves is meant to reveal whether repetition of the same cue reinforces performance for a specific modality and, conversely, whether switching from one cue modality to another makes performance more challenging.

### Sources

All content was generated by humans.

## Results

### Basic features of the urgent multisensory task

We developed the compelled multisensory choice (**CMC**) task to investigate how visual and auditory stimuli are combined during choices that require top-down control (Fig. 1). In each trial, the participant makes a saccadic choice by looking at one of two peripheral targets (gray rings) according to the location of one or two sensory cues. Each trial may be unisensory visual (a spot of light is presented), unisensory auditory (a noise burst is presented), or multisensory (a spot of light and a noise burst are simultaneously presented). The participant is instructed to look either toward or away from a particular stimulus. This setting permits a wide variety of spatial configurations and task rules for exploring how endogenous goals and exogenous factors dictate multisensory performance. Here we consider three task variants. In the first one (CA, Fig. 1a), multisensory trials consist of visual and auditory cues that are presented on the same side and are associated with the same response rule, which is to look away from any stimulus. In the second and third variants, multisensory trials consist of visual and auditory stimuli that are presented on opposite sides and are associated with different sensory-motor rules (saccade away from the visual cue; saccade toward the auditory cue). In the second variant (IVA, Fig. 1b) the visuo-motor rule is applicable in all trials, whereas in the third variant (IAP, Fig. 1c) it is the audio-motor rule which is always applicable.

**Figure 1.**
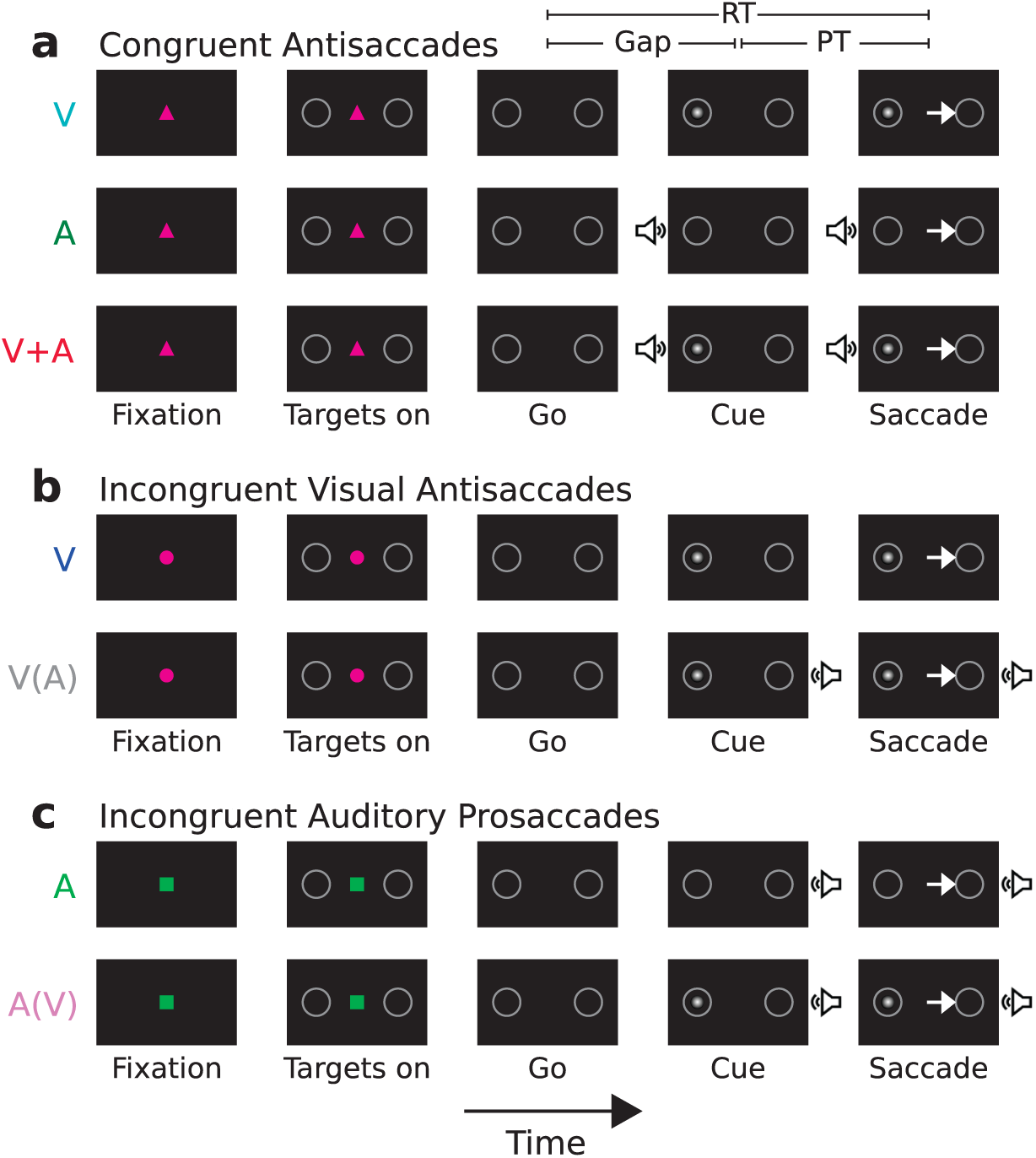
Schematic layout of trials in each variant of the compelled multisensory choice (CMC) task. Following fixation on a central stimulus (Fixation), open targets appear on either side of it (Targets on). After a delay (300–500 ms), the central fixation shape disappears (Go), instructing the participant to respond. Although the correct choice is not known at this point, the participant must strive to initiate an eye movement (Saccade) to one of the targets within a limited reaction time (RT) window (RT. 400 ms). An auditory and/or a visual stimulus is presented (Cue) after the go signal, once a variable time interval has elapsed (Gap, 0–250 ms). The correct choice depends on the cue location and the task rule (an arrow marks the correct saccade in each case). The cue processing time (PT) determines the probability of success in each trial. **a,** Congruent antisaccade (CA) variant. Unisensory visual (V, top row), unisensory auditory (A, middle row), and multisensory audiovisual trials (V+A, bottom row) were interleaved. The fixation shape was a pink triangle, the multisensory cues were presented on the same side, and the goal was to look in the direction opposite to any stimulus, visual and/or auditory. **b**, Incongruent visual antisaccade (IVA) variant. Unisensory visual (V, top row) and multisensory audiovisual trials (V(A), bottom row) were interleaved. The fixation shape was a pink circle, the multisensory stimuli appeared on opposite sides, and the goal was to look in the direction opposite to the visual stimulus. **c**, Incongruent auditory prosaccade (IAP) variant. Unisensory auditory (A, top row) and multisensory audiovisual trials (A(V), bottom row) were interleaved. The fixation shape was a green square, the multisensory cues appeared on opposite sides, and the goal was to look in the direction of the auditory stimulus.

As in other urgent-task designs, in the CMC task time pressure is imposed by first instructing the participant to make a choice (Fig. 1, Go) and then, after a delay (Gap), presenting the relevant stimulus that determines the correct response (Cue). Because the delay is unpredictable (0–250 ms) and the time allowed for responding is limited (RT,:S 400 ms), the participant must hurry or else risks aborting the trial. Over many trials, a wide range of outcomes is produced, going from fast guesses yielding chance performance to fully informed choices approaching 100% correct.

A critical consideration is that the variable that best distinguishes such outcomes is the PT, which corresponds to the period of time during which the sensory cue can be processed before a saccadic response is initiated (Fig. 1a, PT). As such, the psychophysical function that is most useful for characterizing performance in urgent tasks is what we call the “tachometric” curve, which is the curve that results when the fraction of correct choices is plotted as a function of PT (Methods). This psychometric function describes how the likelihood of success changes as time advances over the course of a trial.

Before discussing any multisensory interactions, we consider a condition that serves as a key reference for interpreting all the data — the unisensory visual (V) condition, in which the participant must make an eye movement away from a lone visual stimulus (Fig. 1a, top row). We discuss the V trials from the CA variant, but those from the IVA variant were qualitatively the same (see further below). This trial type corresponds to an urgent, visual antisaccade task.

Antisaccades are characterized by a conflict between the involuntary urge to look at a novel, salient stimulus and the voluntary intent to follow the task instructions and look away (Munoz and Everling, 2004; Coe and Munoz, 2017). Nowhere does this conflict manifest more clearly than in the tachometric curve obtained under urgent conditions (Salinas et al., 2019; Goldstein et al., 2022). As in past antisaccade experiments, the V tachometric curve in the current dataset displays a distinctive shape marked by three phases (Fig. 2). At short PTs (,:S 100 ms) performance hovers around chance (50% correct), indicating that participants simply guess if the cue is not seen for a sufficiently long period of time. At long PTs (;:: 200 ms) performance rises steadily toward an asymptotic performance level close to 100% correct, indicating that participants apply the task rule almost flawlessly when given enough time. And at intermediate PTs (∼100–200 ms) performance dips substantially below chance, indicating that in this range participants are particularly prone to looking at the cue and making errors. Such oculomotor capture results from fast, bottom-up signals that promote reflexive saccades to salient objects (Theeuwes, 2010, 2025). In contrast, the later goal-driven recovery toward asymptotic performance is the consequence of slower, voluntary guidance to the appropriate target (Salinas et al., 2019; Goldstein et al., 2022, 2024; Oor et al., 2023; Kattner et al., 2026).

**Figure 2.**
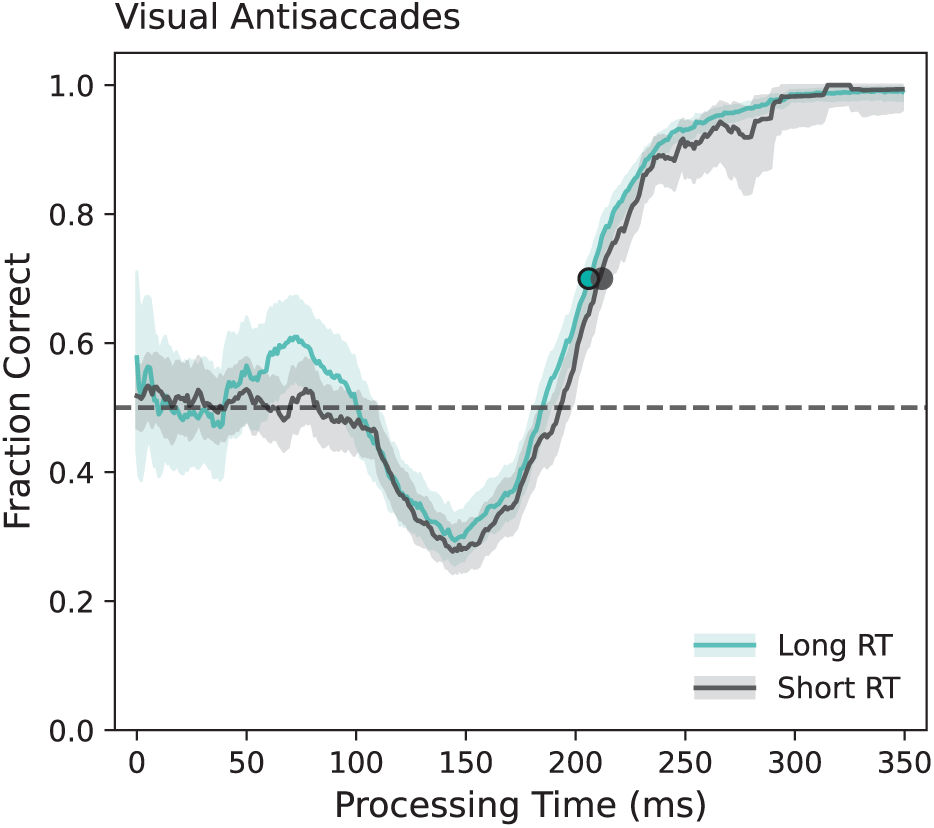
Performance during visual antisaccades. The tachometric curve describes the probability of making a correct response as a function of cue PT, which is measured from cue onset to saccade onset in each trial. Trials are then sorted into PT bins (40 ms width, sliding every 1 ms), and the fraction correct is computed for the data in each bin. Cyan and dark gray curves are based on the slower (long RT) and faster (short RT) halves of the data, respectively, found by splitting the RTs down the median. The RT was measured from the onset of the choice targets (Targets on) to saccade onset. Data are from the unisensory visual (V) condition in the CA variant, with trials pooled over 12 participants. Light shades indicate 95% confidence intervals (CIs) from binomial statistics. Circles mark the rise points of the curves. Horizontal dashed line marks chance performance

The preeminence of the PT is worth stressing. In the current dataset, a lawful relationship exists between RT and accuracy, as would be expected (Wickelgren, 1977; Chittka et al., 2009), but it is rather loose and gives no indication of the conditions that facilitate or prevent cue-driven capture. In contrast, the tachometric curve depicts a relationship between perceptual performance and the time specifically dedicated to cue processing that is largely invariant to motor urgency (Salinas et al., 2014). This can be demonstrated by splitting the data by the median RT and noting that the resulting curves are largely the same (Fig. 2, cyan versus dark gray traces) — even though the two data halves had widely different mean RT and accuracy values (for long RTs: 354 ms and 77% correct; for short RTs: 188 ms and 62% correct).

We used three quantities to characterize each tachometric curve (Methods). The curve rise point is the PT at which performance first exceeds 70% correct (Fig. 2, circles). The probability of capture, or *P_CAP_*, measures the magnitude of the below-chance dip between 0 (for a curve that always stays at or above the chance level) and 1 (for a curve that always stays at or below the chance level), so an intermediate value denotes a curve that swings both below and above chance. Finally, the asymptotic fraction correct is defined as the mean performance over all trials with PTs longer than the curve rise point. For the V curves based on long and short RTs just mentioned (Fig. 2), the values were 206 and 212 ms for the rise point, 0.199 and 0.260 for *P_CAP_*, and 0.94 and 0.93 for the asymptotic fraction correct, and none of these paired values were decidedly different (i.e., the 95% CIs overlapped to some degree; Methods).

With these insights at hand, we now consider how antisaccade performance varies between unisensory and multisensory conditions.

### Multisensory integration during urgent, congruent antisaccades

The CA variant of the task (Fig. 1a) was designed as a standard multisensory experiment in which a stimulus must be detected and located, except that participants were meant to look *away* from it. The question was whether during goal-driven choices the presence of two multisensory cues would manifest as if a single, stronger, integrated cue had been shown. Participants made anti-saccades guided either by a unisensory auditory (A) cue, by a unisensory visual (V) cue, or by a multisensory (V+A) cue for which the auditory and visual stimuli were presented simultaneously on the same side. That is, the multisensory condition was both stimulus-and target-congruent. The three trial types were randomly interleaved, and a tachometric curve was generated for each type (Fig. 3).

**Figure 3.**
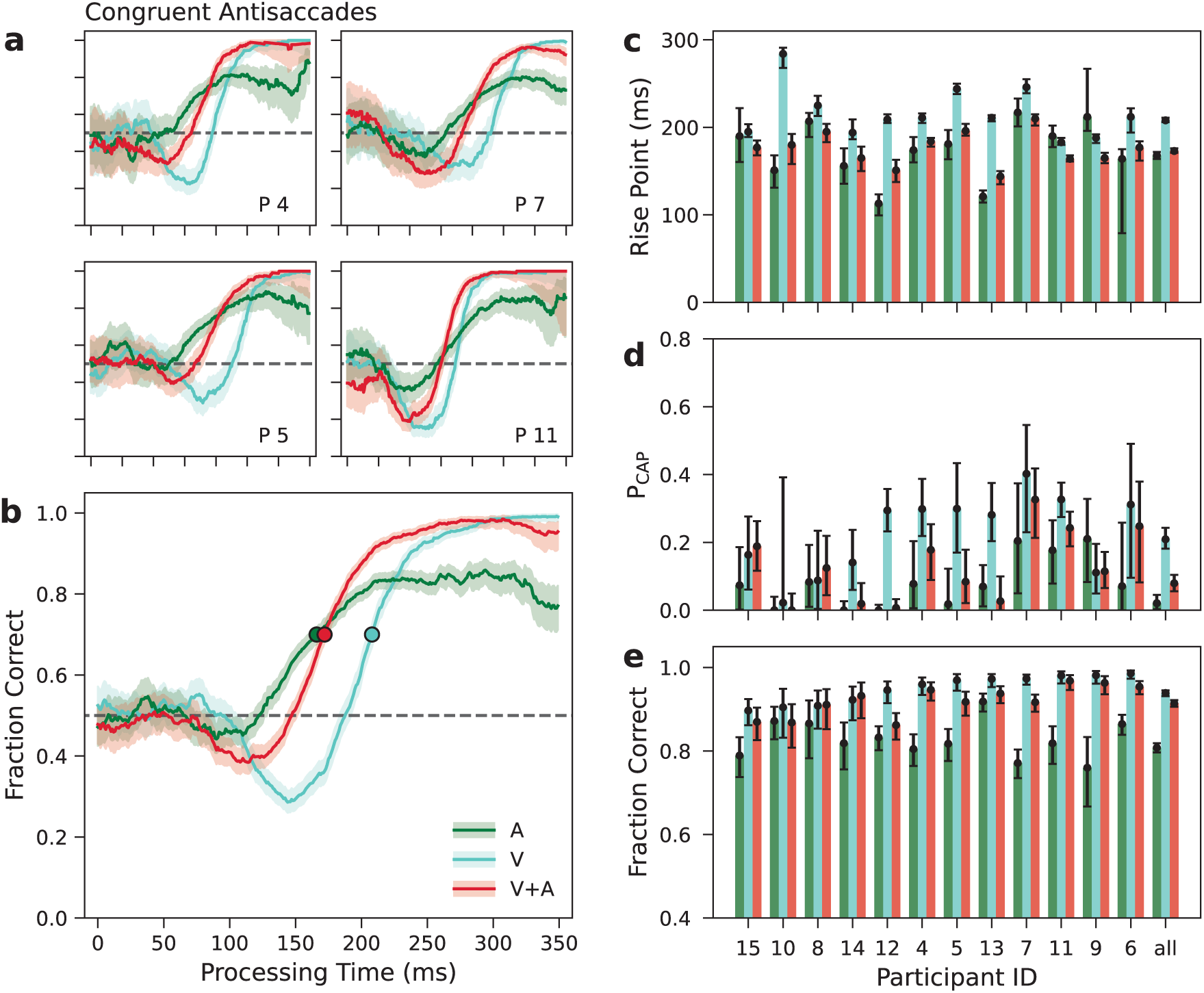
Performance during congruent antisaccades (CA). **a, b**, Tachometric curves for four example participants (**a**) and for the data pooled over participants (**b**). Traces are for auditory alone (A, green), visual alone (V, cyan), and congruent multisensory trials (V+A, red). Light shades indicate 95% CIs from binomial statistics. Circles indicate rise points. Horizontal dashed lines mark chance performance. c, Rise points of the tachometric curves in the A (green bars), V (cyan bars), and V+A (red bars) conditions for each participant. Errorbars indicate 95% CIs from bootstrap. **d**, As in c but for the probability of capture (PCAP). **e**, As in **c** but for the average fraction correct in the asymptotic range (PT > rise point). Errorbars indicate 95% CIs from binomial statistics. Participants are sorted based on the asymptotic fraction correct in the V condition (cyan bars). Results from pooled data are labeled as ‘all’

As previously mentioned, the V tachometric curve (Fig. 3a, b, cyan traces) demonstrated (1) an early dip below chance consistent with an involuntary, stimulus-driven signal that favors saccades to the cue, and (2) a later rise toward 100% correct consistent with a voluntary, goal-driven signal that favors (anti) saccades to the desired target. The tachometric curve for auditory anti-saccades (A; Fig. 3a, b, green traces) differed from the V curve (cyan traces) in three ways: it rose considerably earlier, demonstrated a minimal dip, and saturated at a lower ceiling.

The faster rise is not surprising given that the latencies of auditory stimuli in the cortex are shorter than those of visual stimuli (Schroeder et al., 1998; Bair et al., 2002; Camalier et al., 2012; Nourski et al., 2014), which accounts for correspondingly faster RTs in detection tasks (Elliott, 1968; Welford, 1980; Shaw et al., 2020). Our CA data demonstrate that similar timing differences between vision and audition manifest in perceptual processing speed independently of RT effects. When pooling the data across participants, the rise point for A trials (166 ms in [165, 171], 95% CI) was lower than that for V trials (208 ms in [206, 211]) by 42 ms (Fig. 3b, green and cyan circles). This difference was consistent across individual participants (Fig. 3c, green and cyan bars); for the majority of them (8 of 12) the A rise point was decidedly lower than the V rise point (i.e., no overlap between 95% CIs; Methods).

The two other differences between V and A trials are notable given the low intensity of the visual cue and high intensity of the auditory (Methods). In spite of its loudness, the auditory stimulus produced very weak capture (Fig. 3d, green bars). For the pooled data, *P_CAP_* in A trials was just slightly above zero (0.038 in [0.025, 0.057], *P* (0) *<* 0.0001; Methods). Likewise, *P_CAP_* was positive with high significance (*P* (0) *<* 0.001) only for 3 of 12 individual participants (for example, for participants 7 and 11 in Fig. 3a). In contrast, in spite of its low luminance, the visual stimulus produced consistent capture (Fig. 3d, cyan bars). In V trials *P_CAP_* was 0.224 (in [0.196, 0.249], *P* (0) *<* 0.0001) for the pooled data, a value well above zero. Likewise, individual values were positive with high significance (*P* (0) *<* 0.001) for 10 of 12 participants. The differences in performance at long PTs were similarly distinct (Fig. 3e, green and cyan bars). For the pooled data, the asymptotic fraction correct for A trials (0.81 in [0.80, 0.82]) was well below that for V trials (0.94 in [0.93, 0.95]), and the difference went in the same direction for all 12 participants.

These results reveal a substantial functional asymmetry between visual and auditory modalities that is consistent with the idea that, for certain behaviors, the former dominates over the latter (Colavita, 1974; Spence et al., 2012; Li et al., 2017; Malevich et al., 2026). When an antisaccade is requested, the bottom-up signal associated with a weak (low-salience) visual cue exerts stronger exogenous capture than that associated with a loud auditory cue — and yet, the same visual cue also leads to a more reliable endogenous signal for shifting the eyes to the correct target. The visual cue seems to have a stronger impact on target selection, for better or worse.

What was the consequence of presenting the visual and auditory cues simultaneously? In general, performance in the V+A condition (Fig. 3, red bars and traces) reflected the relative advantages of the two cues, i.e., the lower latency of the auditory and higher reliability of the visual. Compared to performance in the V condition, the added presence of the auditory cue decreased the magnitude of exogenous capture and sped up the goal-driven rise with little impact on asymptotic accuracy (Fig. 3b, compare red and cyan traces). For the pooled V+A trials, *P_CAP_* was intermediate between those measured in A and V trials (0.095 in [0.075, 0.103], *P* (0) *<* 0.0001), but was closer to the former (0.038) than the latter (0.224). The rise point (172 ms in [170, 173]) was just above that for A trials (166 ms; Fig. 3b, green and red circles). And the asymptotic fraction correct (0.91 in [0.90, 0.92]) was just below that measured during V trials (0.94). These same patterns were common in the data from individual participants (Fig. 3a).

### A statistical benchmark for assessing the multisensory benefit

Clearly, the multisensory product evolves as a function of PT, and the evolution is somewhat complicated (Fig. 3b, red trace). For instance, compare the multisensory choices that were predominantly stimulus driven (i.e., with PT,:S 172 ms) with those that were predominantly goal-driven (i.e., with PT;:: 172 ms). For the former, performance in the V+A multisensory trials was generally intermediate between that in the A and V conditions; for the latter, the multisensory performance was generally better than for either of the unisensory conditions.

To attain a more intuitive understanding of these temporal changes, we developed a statistical benchmark for the integration process that fits the conditions of the CMC task (Methods). During sensory-driven choices, performance given two informative signals may be more or less accurate than with just one, depending on how such signals interact as they drive the intended behavior. The concept of conditional independence is useful for characterizing such interaction probabilistically (Coen et al., 2023; Salinas and Stanford, 2024; Oor et al., 2025). When the A and V signals are conditionally independent of each other relative to choice outcome, it is possible to make a parameter-free, quantitative prediction about their joint effect on performance (i.e., on the probability of making a correct choice) based on how each signal relates to behavior individually (see Methods for details). In practice, this means that a prediction for the multisensory tachometric curve can be generated using the two unisensory curves.

In general, the data from the CA experiment were qualitatively close to the statistical predictions derived by assuming conditional independence (Fig. 4a, b). Relative to the V case, the expected multisensory curves showed an attenuated dip below chance and an earler rise toward asymptotic accuracy, just as seen experimentally. In assessing these qualitative trends, it is useful to keep in mind that the conditional independence criterion considers the accuracy of the unisensory choices with respect to chance (Equation 5). If the A and V cues both yield above-chance performance, then the predicted multisensory performance is above both unisensory values (agreement between cues reinforces their common trend); likewise, if both cues yield below-chance performance, the prediction is lower than both unisensory values; however, if the probability correct is above chance for one cue but below chance for the other, their predicted combination is an intermediate probability (a compromise is struck between conflicting effects). The experimental data are clearly consistent with these distinctions drawn in relation to chance performance (Fig. 3b).

**Figure 4.**
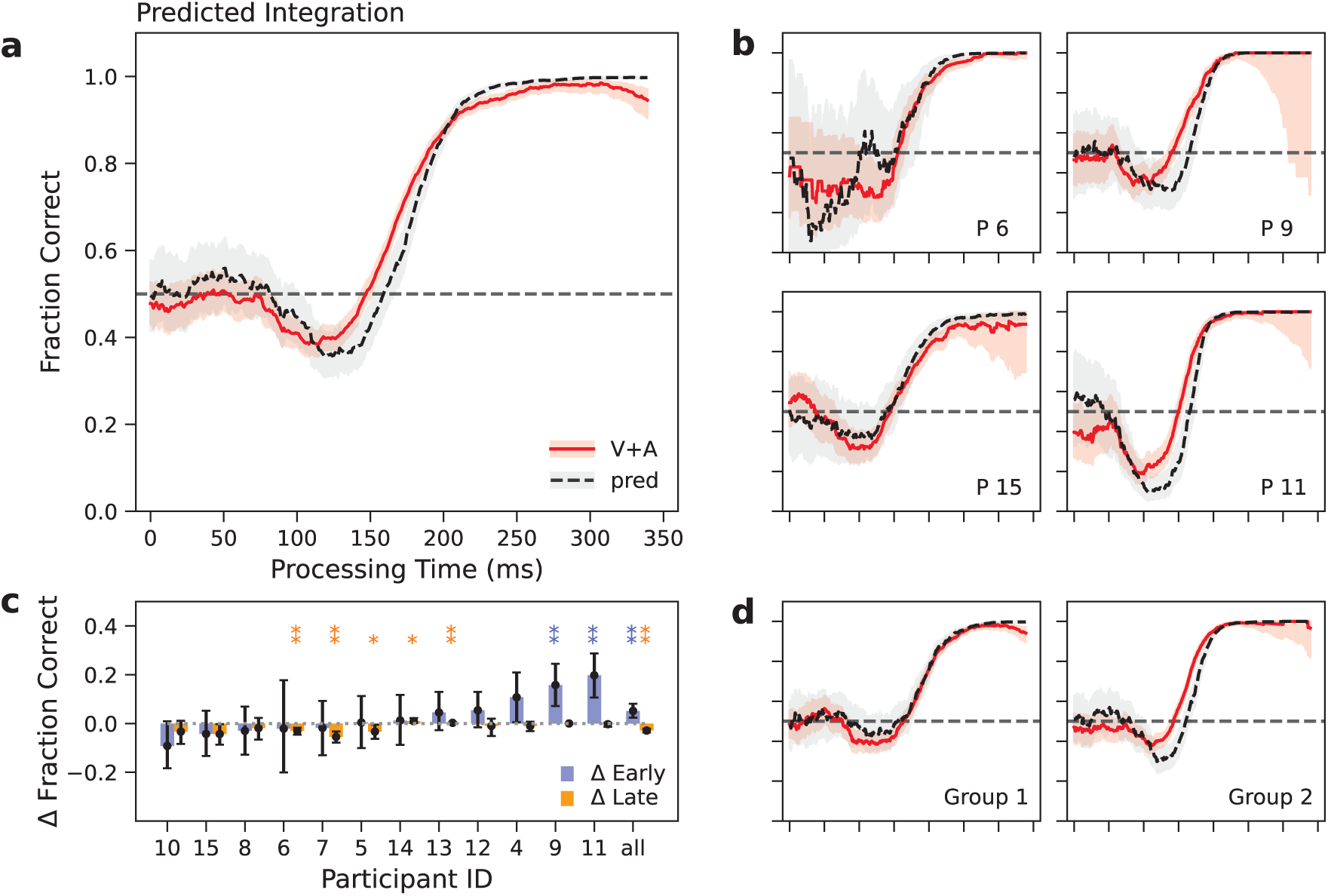
Test of multisensory integration in CA trials. **a, b**, Red lines show tachometric curves from multisensory trials (V+A). Black dashed lines show the predicted tachometric curves (pred) based on conditionally independent contributions from the two unisensory modalities (V and A trials). Results are for the data pooled over participants (**a**) and for four example participants (**b**). Light shades indicate 95% CIs from binomial statistics. Horizontal dashed lines mark chance level. **c**, Difference between observed and predicted fraction correct in an early-rise range (100–208 ms, purple bars) and in a late-rise range (208–350 ms, orange bars) of PTs for each participant. Errorbars indicate 95% CIs from bootstrap. Participants are sorted in order of increasing difference (empirical minus predicted) in the early range. Results from pooled data are labeled as ‘all’. Stars mark significance (one star, p < 0.01; two stars, p < 0.001) for values that were statistically different from zero. **d**, Pooled tachometric curves for two groups of participants that had the smallest (left panel, n = 6) or the largest (right panel, n = 6) Δ Early. Same format as in a.

Quantitatively, the empirical curves demonstrated small but significant discrepancies with respect to the predictions. During the early rise in performance (100 ≤ PT ≤ 208 ms) the observed accuracy was slightly above the prediction, whereas during the later part of the rise (208 ≤ PT ≤ 350 ms) it was slightly below (Fig. 4a). In terms of average fraction correct within each of these PT ranges, both differences were highly significant for the pooled data (*p* = 0.0003 for the early rise, *p <* 0.0001 for the late rise, from resampling tests; Methods). Thus, on average, multisensory performance for early goal-driven choices was slightly above that expected from conditionally independent A and V signals (Fig. 4c, purple bars). In contrast, for late goal-driven choices that approach ceiling accuracy, performance was slightly below the expectation (Fig. 4c, orange bars).

Interestingly, though, these two effects in the pooled data, for early and late PT ranges, seemed to be due primarily to contributions from different participants (Fig. 4c, d). Our sample could be divided into two groups. For one group (participants 5, 6, 7, 8, 10, 15), performance in V+A trials was slightly below the prediction during the early rise (*p* = 0.045 for the data pooled over the group) and definitely below the prediction during the late rise (*p <* 0.0001). For the other group (participants 4, 9, 11, 12, 13, 14), performance was higher than the prediction during the early rise (*p <* 0.0001) and no different from the prediction during the late rise (*p* = 0.23). Pooling the data separately for these two groups produced contrasting profiles (Fig. 4d), and these were more representative of the results from individual participants (Fig. 4b, note different patterns for P6 and P15 versus P9 and P11) than the overall average (Fig. 4a). In conclusion, some participants integrated the multisensory cues more effectively than others; however, the divisions were somewhat obscure because successful integration (performance above expectation) was most evident in the early window whereas suboptimal integration (performance below expectation) was most evident in the late window.

In summary, the CA experiment revealed a form of multisensory integration that varied rapidly over time, first combining predominantly stimulus-driven signals and then transitioning to predominantly goal-driven ones. The integration was qualitatively different before and after the transition, but the entire timecourse was highly consistent with a statistical prediction (based on the assumption that the unisensory cues are conditionally independent with respect to choice outcome). There was some variability to this process across participants; some exceeded the statistical benchmark whereas others fell short. Nevertheless, relative to either cue alone, in this case multisensory cues yielded a clear performance benefit during goal-driven choices.

### Urgent multisensory integration with incongruent cues

The IVA and IAP variants of the task (Fig. 1b, c) were designed as complementary parts to a multisensory experiment in which, during multisensory trials, the two cues were spatially incongruent, i.e., they were presented on diametrically opposite locations. Critically, though, the spatial relationship never changed and the response rules associated with such cues never instructed conflicting saccadic choices. Unisensory trials required participants to make either antisaccades away from a V cue, in IVA blocks, or prosaccades toward an A cue, in IAP blocks. During IVA multisensory trials the additional auditory cue could, in principle, enhance the visually driven anti choice by guiding the saccade early and directly toward the correct target (via an implicit pro rule). Likewise, during IAP multisensory trials the additional visual cue could, in principle, reinforce the sound-driven pro choice by confirming the correct target location (via an implicit anti rule). Thus, while multisensory trials were stimulus-incongruent due to spatial misalignment of the cues, the consistent application of different cue-response rules would have kept them target-congruent. The question was whether, in either variant, the participants would be able to exploit such consistency and benefit from the presence of two cues instead of one.

What are the possible scenarios for the outcome of this experiment? If integration is at all feasible, one possibility is that a performance benefit is eventually seen at long PTs, as in the CA experiment. This would be in line with the effects of practice and repetition on attentional selection (Jiang and Sisk, 2019), and with numerous experiments in which crossmodal features that seemed to be unrelated, or related only in an abstract way, turned out to be tightly bound perceptually (e.g., auditory pitch and visual size; Laurienti et al., 2004; Ludwig et al., 2011; Spence, 2011). On the other hand, given that it is generally difficult to perform multiple sensory-motor tasks simultaneously (Pashler, 1994; Marois and Ivanoff, 2005), an alternative is that participants predominantly track one stimulus in each trial, essentially filtering out the other one. If participants focused on the stimulus that is persistent in each task variant, this would predict multisensory performance that is very similar to that in the corresponding unisensory trial type. As we describe next, results were close to the latter scenario, but with the caveat that stimulus-driven interactions were still clearly visible.

In the IVA condition (Fig. 5), tachometric curves were generally similar for unisensory (V) and multisensory (V(A)) trials, particularly during the predominantly goal-driven rise in performance (Fig. 5a, b). For the pooled data the rise points in V (226 ms in [224, 230]) and V(A) trials (222 ms in [220, 224]) differed by only 4 ms, and for individual participants the values were generally very similar as well; these were decidedly different only in 3 cases (Fig. 5a, c, P8, P12, P14). The asymptotic fraction-correct values were separated by only 0.02 in the pooled datasets, and were decidedly different for only 1 of the participants (Fig. 5e). The largest contrast between IVA trial types was seen in *P_CAP_*, which was 0.289 (in [0.258, 0.323], *P* (0) *<* 0.0001) for the pooled V data versus 0.071 (in [0.040, 0.105], *P* (0) *<* 0.0001) for the pooled V(A) data, with the difference going in the same direction for 9 of 13 participants (Fig. 5d). These results indicate that, in the IVA variant of the task, multisensory performance was better than unisensory only for short-PT trials (∼50–210 ms), during which the bottom-up signals influenced the choice most directly. In that range, the auditory stimulus acted as if to pull the saccade toward it — though not enough to completely overcome the draw of the visual cue. Later, during choices that were predominantly goal-driven, the auditory cue had hardly any impact.

**Figure 5.**
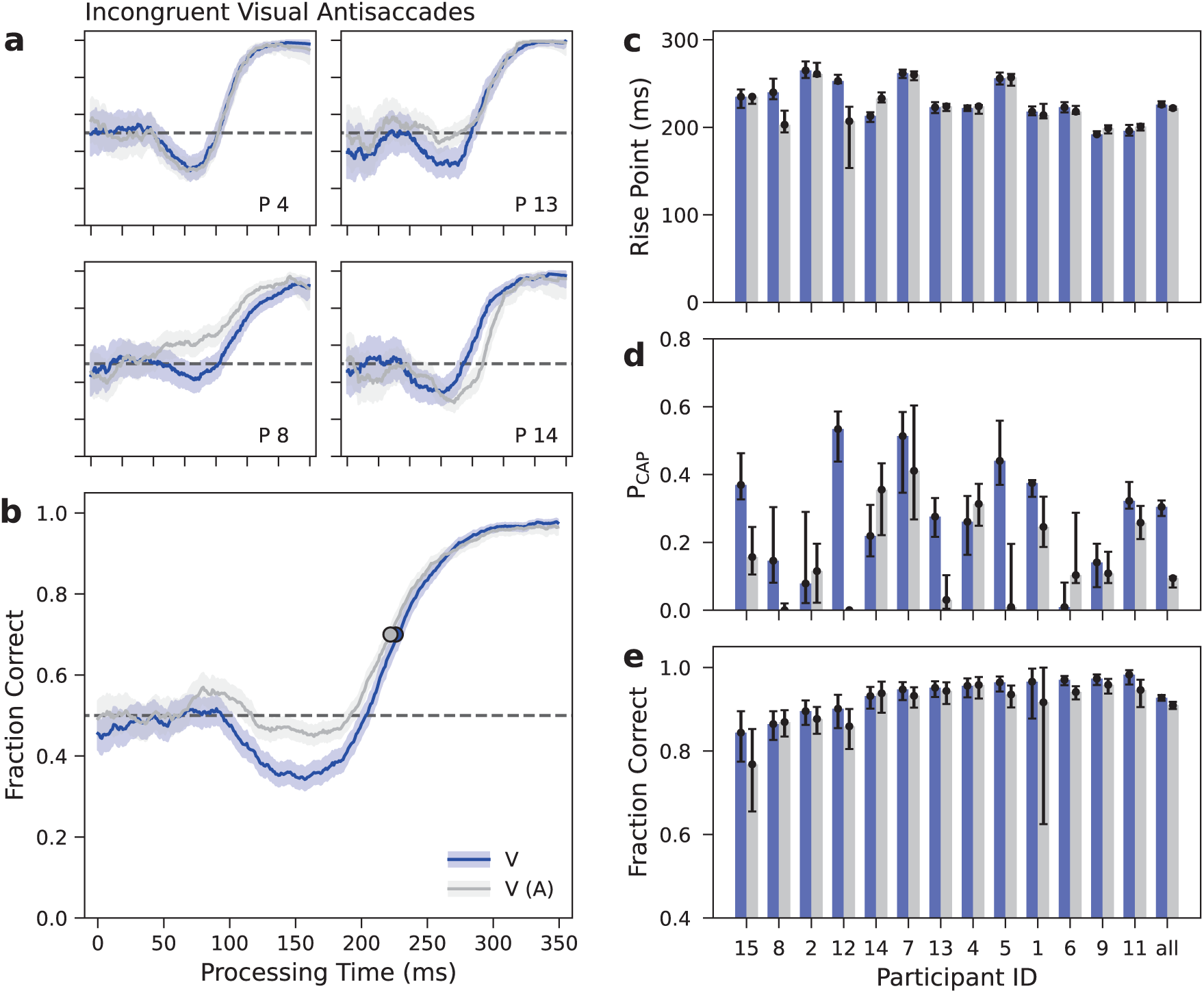
Performance during incongruent visual antisaccades (IVA). **a, b**, Tachometric curves for four example participants (**a**) and for the data pooled over participants (**b**). Traces are for unisensory visual (V, blue) and incongruent multisensory trials (V(A), gray). Light shades indicate 95% CIs from binomial statistics. Circles indicate rise points. Horizontal dashed lines mark chance performance. **c**, Rise points of the tachometric curves in the V (blue bars) and V(A) conditions (gray bars) for each participant. Errorbars indicate 95% CIs from bootstrap. **d**, As in c but for the probability of capture (PCAP). **e**, As in **c** but for the average fraction correct in the asymptotic range (PT > rise point). Errorbars indicate 95% CIs from binomial statistics. Participants are sorted based on the asymptotic fraction correct in the V condition (blue bars). Results from pooled data are labeled as ‘all’.

In the IAP condition (Fig. 6), the auditory cue instructed a prosaccade and tachometric curves generally rose much sooner, consistent with an early, bottom-up auditory signal driving the choice. In this case, tachometric curves were nearly identical for unisensory (A) and multisensory (A(V)) trials during the early rise in performance (Fig. 6a, b). The A and A(V) rise points typically occurred at 78 ms (in [76, 79], pooled data) and 77 ms (in [75, 79]), respectively, and were not decidedly different for any of the participants (Fig. 6c). Moreover, consistent with the universal rise in performance seen before and past each rise point, all but two of the *P_CAP_* values were either zero or far from significant (Fig. 6d; *P* (0) *>* 0.3 for all participants except P10 in A(V) trials and P6 in A trials, for which *P* (0) = 0.05 and 0.03, respectively). Notably, however, the A and A(V) curves clearly differed for PTs longer than ∼100 ms — the point at which the visual cue starts manifesting via captured saccades during V trials (Fig. 3b, cyan trace; Fig. 5b, blue trace). For the pooled data, the asymptotic fraction correct was 0.95 in A trials and 0.90 in A(V) trials (both with uncertainties *<* 0.01), with the difference going in the same direction for 12 of the 13 participants (Fig. 6e). The incongruent visual cue clearly hindered performance during auditory prosaccades.

**Figure 6.**
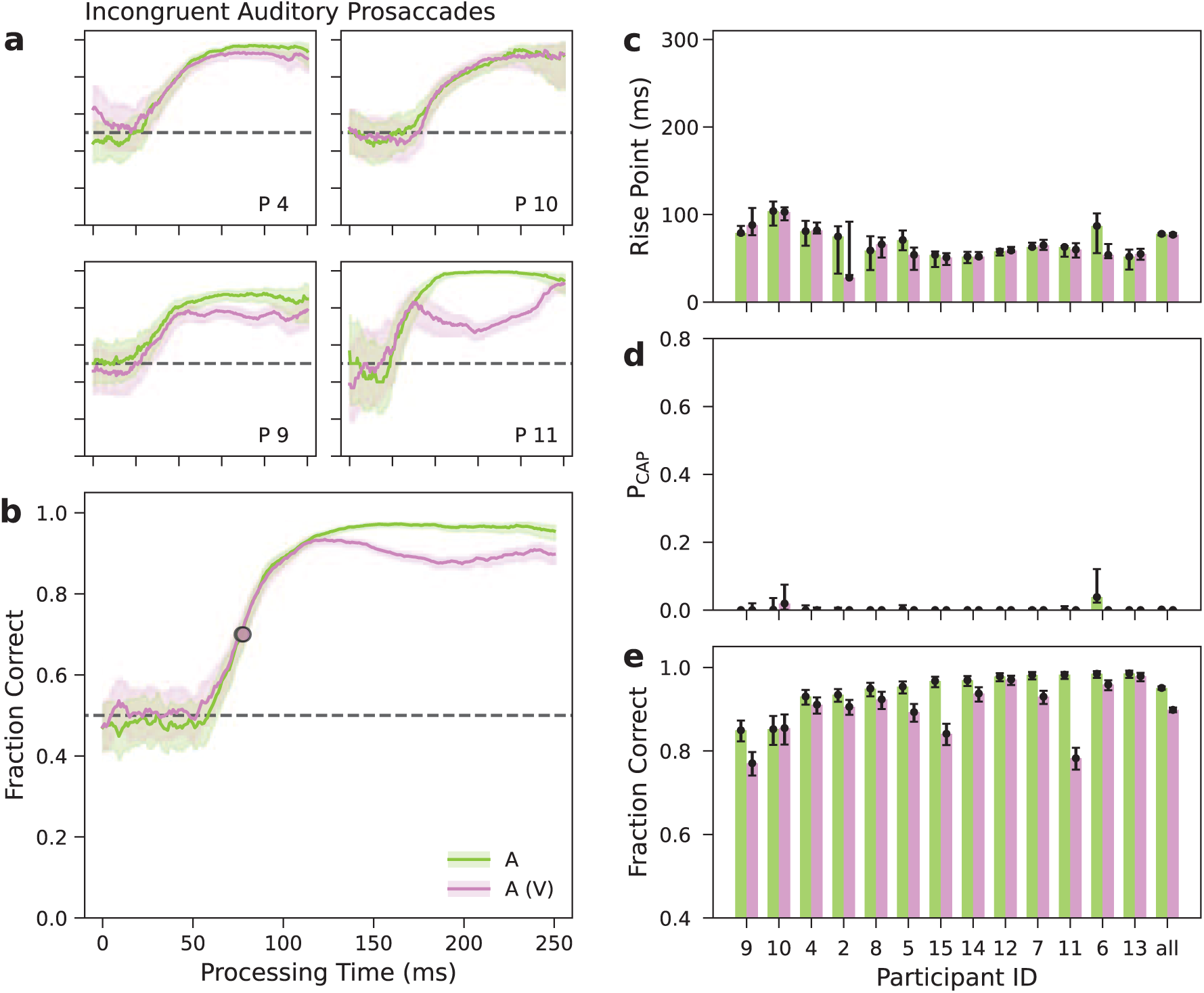
Performance during incongruent auditory prosaccades (IAP). **a**, **b**, Tachometric curves, which plot the probability of making a correct response as a function of processing time, for four example participants (**a**) and for the data pooled over participants (**b**). Traces are for unisensory auditory (A, green) and incongruent multisensory trials (A(V), pink). Light shades indicate 95% CIs from binomial statistics. Circles indicate rise points. Horizontal dashed lines mark chance performance. **c**, Rise points of the tachometric curves in the A (green bars) and A(V) conditions (pink bars) for each participant. Errorbars indicate 95% CIs from bootstrap. **d**, As in **c** but for the probability of capture (*P_CAP_*). **e**, As in **c** but for the average frac-tion correct in the asymptotic range (PT *>* rise point). Errorbars indicate 95% CIs from binomial statistics. Participants are sorted based on the asymptotic fraction correct in the A condition (green bars). Results from pooled data are labeled as ‘all’.

Although the shapes of the respective tachometric curves were very different, the results for the IVA and IAP variants were, in fact, highly consistent with each other. Broadly speaking, in both cases the performance profile with multisensory cues was close to that with the persistent unisensory cue. And examining the distinctions more closely, in both cases the additional cue biased performance as if the saccades were reflexively drawn toward it (due to its salience). This interpretation reconciles the opposite effects on timing and accuracy observed in the two variants. During IVA trials, the short-latency auditory cue acted early (relative to the V rise in performance) to increase the fraction of correct choices. In contrast, during IAP trials, the longer-latency visual cue acted late (relative to the A rise in performance) to decrease the fraction of correct choices. Given their timing and direction, these competitive effects are consistent with exogenous or stimulus-driven capture (Theeuwes, 2010, 2025); it just so happened that in the IVA variant the location of the additional auditory cue was aligned with the correct V(A) choice, whereas in the IAP variant the location of the additional visual cue was aligned with the incorrect A(V) choice.

### Incongruent conditions deviate from integration predictions

Taken together, the results from the IVA and IAP variants contrast dramatically with those from the CA variant, in which the two cues were associated with the same stimulus-response rule and performance was close to that predicted by the conditional-independence benchmark (Fig. 4). This can be more readily appreciated by computing an analogous prediction for the IVA/IAP variants. In those variants, A and V trials were never interleaved in the same block, but their corresponding tachometric curves can nonetheless be combined to generate a prediction for the performance to be expected for signals that are integrated when presented together (Fig. 7a). The predicted curve (black dashed trace) represents what is theoretically possible, i.e., what would happen in this case if participants were able to combine the information from the two cues (and their corresponding response rules) with the same efficiency as in the CA case, i.e., such that conditional independence was approximately satisfied.

**Figure 7.**
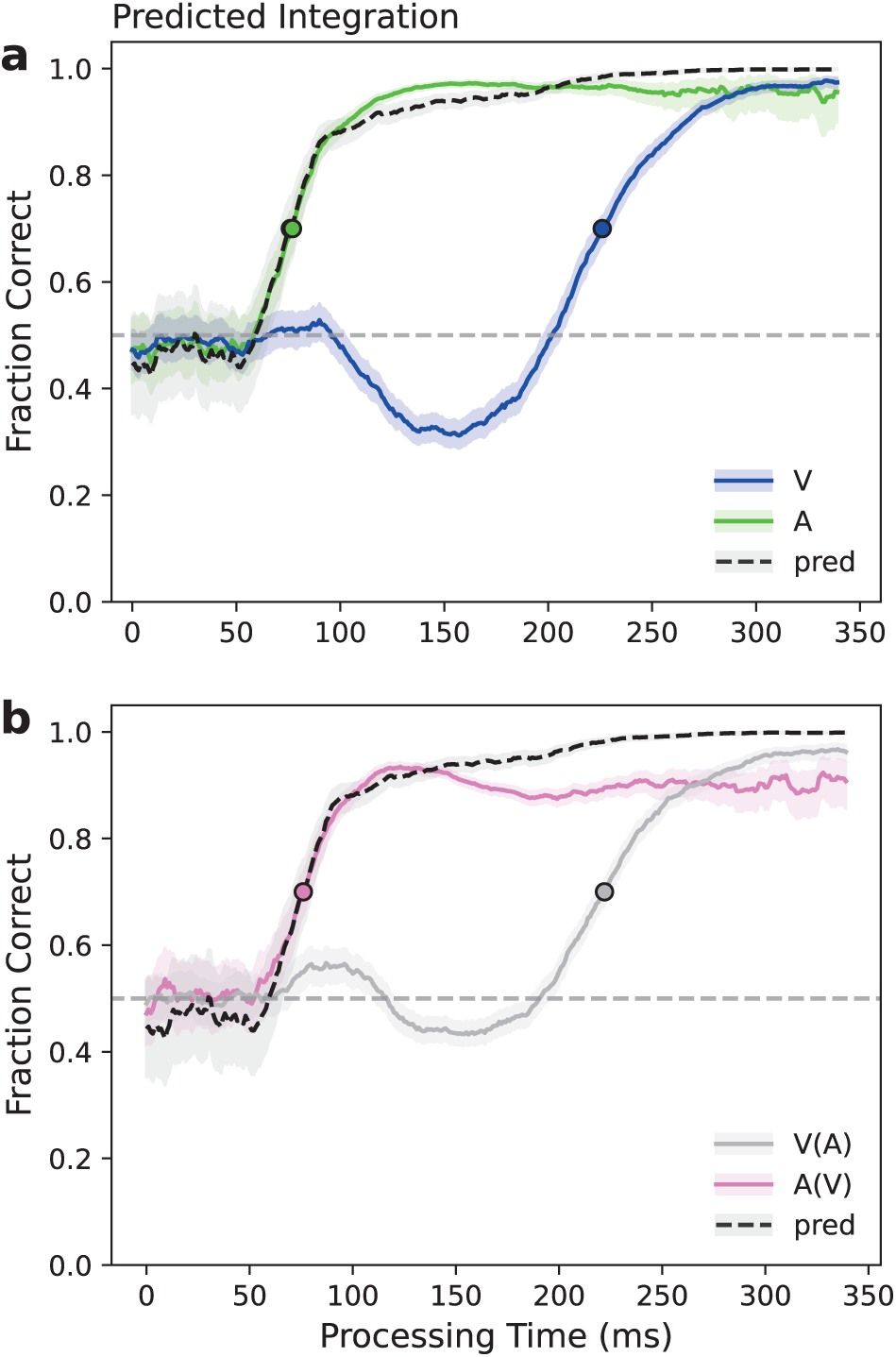
Test of multisensory integration for stimulus-incongruent but target-congruent trials. **a**, Multisensory performance predicted from unisensory trials. Blue and green lines show unisensory tachometric curves for V and A trials from the IVA and IAP variants, respectively (same pooled unisensory curves as in Figs. 5b and 6b). The black dashed line shows the multisensory tachometric curve predicted from the two unisensory curves by assuming conditional independence between the auditory and visual cues with respect to choice outcome. Light shades indicate 95% CIs from binomial statistics. Circles indicate rise points. Horizontal dashed line marks chance level. **b**, Predicted versus observed multisensory performance. Gray and pink lines show multisensory tachometric curves for V(A) and A(V) trials from the IVA and IAP variants, respectively (same pooled multisensory curves as in Figs. 5b and 6b). The black dashed line shows the predicted multisensory tachometric curve (same as in **a**). Neither experimental curve matches the multisensory prediction.

**Figure 8.**
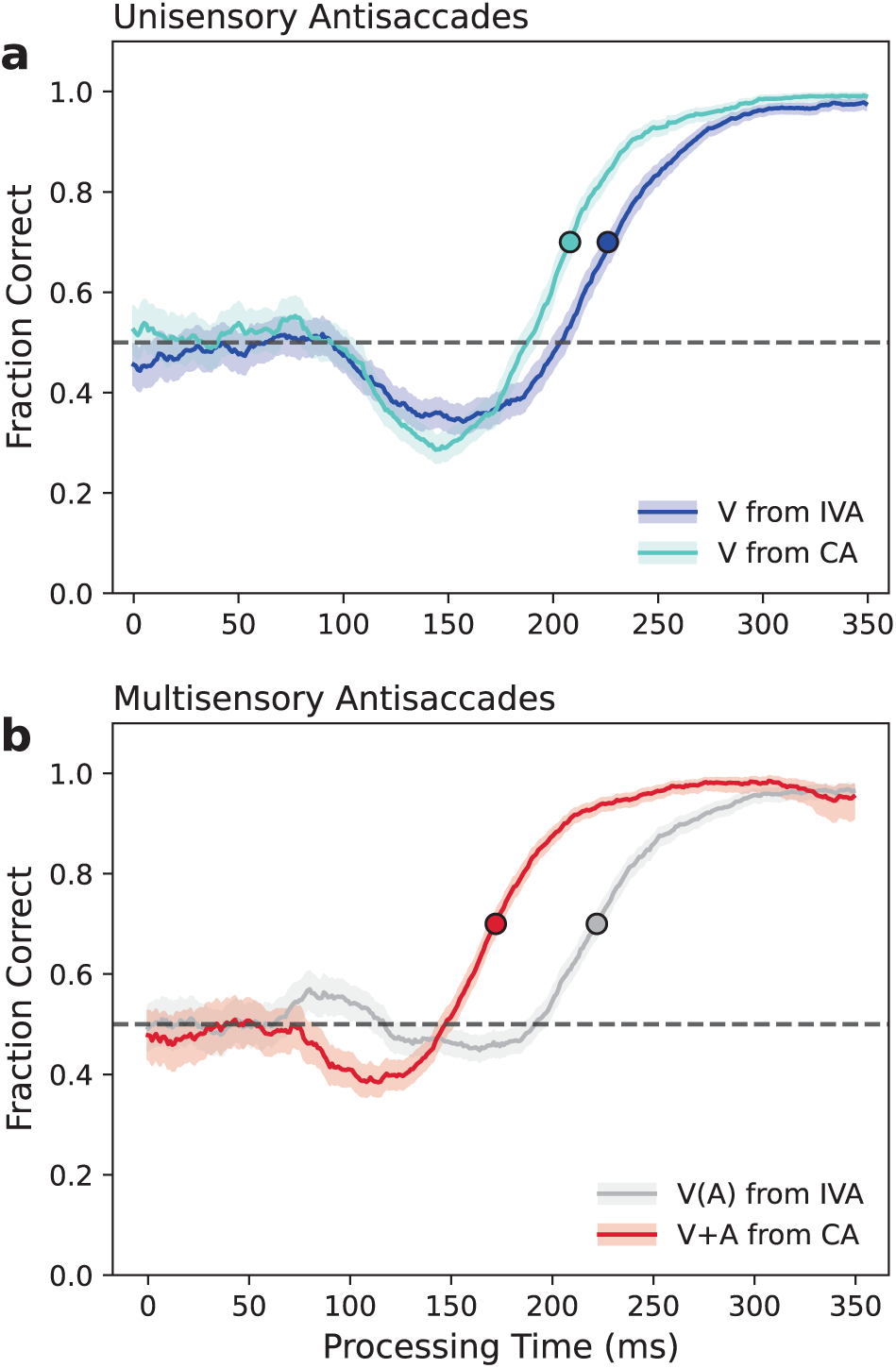
Context-dependence of visual and multisensory performance. **a**, Tachometric curves for unisensory visual trials in the CA (cyan) and IVA (dark blue) contexts. Same pooled traces as in Figs. 3b and 5b, V conditions. Light shades indicate 95% CIs from binomial statistics. Circles indicate rise points. Horizontal dashed lines mark chance performance. Identical visual stimuli directed antisaccades in the two conditions. **b**, Tachometric curves for multisensory trials in the CA (red) and IVA (gray) contexts. Same pooled traces as in Figs. 3b and 5b, V+A and V(A) conditions, respectively. Multisensory performance is drastically different for congruent versus incongruent cues.

Again, when one unisensory signal yields performance above and the other yields performance below chance, the predicted curve strikes a compromise, and otherwise it stipulates that performance should be more extreme than with either signal alone (Fig. 7a). But in this case, for both the V(A) and A(V) multisensory conditions, the observed fraction correct was well below the expectation (Fig. 7b). These comparisons support the conclusions from the preceding subsection: multisensory performance in the IVA and IAP variants was nowhere near what should have been seen if participants had been able to integrate the two spatially disparate cues.

Why? Was effective integration precluded by splitting the V, A, and multisensory conditions into separate experiments (IVA first, IAP second), perhaps? An alternative design would have been to interleave all the incongruent trial types (A prosaccades, V antisaccades, multisensory) within the same blocks of trials, in analogy with the CA experiment. This would have required that participants switch between pro and anti rules from one unisensory trial to another, and would have made the rule for multisensory trials ambiguous. The current design was simpler, in that only one rule had to be actively kept in mind in any block, but in principle it still allowed participants to benefit from the additional, consistently related information presented during multisensory trials.

Still, it could be argued that, in the IVA variant, participants demonstrated minimal integration (Fig. 7b, gray versus black traces) precisely because they had not explicitly practiced the auditory pro rule, or did not realize that they could apply it during multisensory trials. We think this is unlikely, though. In terms of both success rate and processing time costs, making prosaccades is much easier than making antisaccades (Goldstein et al., 2022; Oor et al., 2023; Kattner et al., 2026). This is plainly demonstrated by the comparison between the A antisaccade trials from the CA variant (Fig. 3a, b, green traces) and the A prosaccade trials from the IAP variant (Fig. 6a, b, green traces), which differed markedly in both rise point (by 88 ms) and asymptotic fraction correct (by 0.14). For this reason, we believe that if it had been at all possible for participants to produce multisensory choices predominantly driven by auditory prosaccades in the IVA variant, they would have done so, even without having invoked the auditory pro rule explicitly beforehand. In the case of the IAP variant, participants did have extensive prior practice with visual antisaccades (during the earlier IVA blocks), but the visual cue clearly had no benefit either (Fig. 7, pink versus green and black traces). With this in mind, the two sets of results suggest that spatial incongruence precluded integration and led to cross-modal competition.

### Contextual effects are consistent with attention mechanisms

The filtering and prioritization of sensory information is precisely what defines selective attention (Carrasco, 2011; Moore and Zirnsak, 2017; Xia et al., 2024). With this in mind, note the striking difference of 145 ms between the rise points of the tachometric curves in the V(A) and A(V) conditions (Fig. 7b, gray versus pink traces). What varied between these two multisensory curves was the context, i.e., the surrounding unisensory trials and the instructions given to the participants. However, the stimuli and associated saccade targets were identical in the two cases. The result is a perfect illustration of selective attention at work, with both history-driven and goal-driven mechanisms potentially contributing (Spence et al., 2001; Kattner et al., 2026). Differences in the participants’ internal perceptual (i.e., stimulus-related) and/or cognitive (i.e., instruction-related) settings yielded unequivocally distinct performance profiles over time. Notably, the distinction fades away eventually, for PT;:: 250 ms. Had the paradigm failed to probe earlier PTs, only a minimal residual difference in ceiling accuracy would have been detected, if anything.

Our data demonstrated another contextual effect that is also consistent with the operation of attentional selection during task performance. This is the contrast between visual antisaccades in the CA and IVA variants (Fig. 8a). For the two sets of V trials, the stimuli, targets, and instructions were identical; only the surrounding multisensory trials distinguished the two conditions. And yet, in the CA case performance rose 18 ms earlier than in the IVA case (pooled data; corresponding rise points were 208 ms in [206, 211] and 226 ms in [224, 230]; Fig. 8a, circles). This effect was robust: the rise points were decidedly different for 8 of 11 participants, and the difference went in the same direction for all of them. Differences in *P_CAP_* and asymptotic fraction correct were more modest but also highly consistent across the population. It is clear that the participants’ attentional set differed reliably between the two task variants.

Finally, it is also instructive to ponder how attentional allocation may have differed between the two multisensory conditions from the CA and IVA variants (Fig. 8b). This is a direct comparison of performance in congruent (V+A, red trace) versus incongruent (V(A), gray trace) multisensory trials, and it relates to the question, how exactly did the execution of visually guided antisaccades change depending on the location of the ancillary auditory cue? The influence of such cue began to manifest around the PT = 75 ms mark in both cases (see also Fig. 6b, c), but the most prominent difference was that the congruent tachometric curve rose about 50 ms sooner than the incongruent (rise points were 172 ms in [170, 173] and 222 ms in [220, 224], respectively; Fig. 8b, circles). This can be understood in terms of different patterns of shifts in spatial attention. In the congruent case, as with regular V antisaccades, attention shifts twice: it is first drawn exogenously to the one cue location, sometimes yielding erroneous captured saccades, and then shifts endogenously to the correct target location once the cue information has been interpreted according to the ‘anti’ task rule. Modeling and neurophysiological results support this scenario (Salinas et al., 2019; Oor et al., 2023; Zhu et al., 2024). By contrast, in the incongruent case attention seems to shift three times: it is first drawn exogenously to the auditory cue, then to the longer-latency visual cue, also exogenously, and it is finally redirected back to the target location (i.e., to the auditory cue) via endogenous guidance. The additional shift — or, equivalently, the split of attentional resources — consumes 50 ms of extra PT. This account is consistent with the subtle yet visible tri-phasic shape of the V(A) curve, which first increases above chance, then dips slightly below chance, and finally rises again toward its ceiling (Fig. 8b, gray trace). It is also consistent with a previous study in which similar shifts of attention occured between two diametrically opposite visual stimuli, which led to even larger tri-phasic fluctuations in performance (Goldstein et al., 2024).

### Potential contribution of selection history effects

An alternative mechanism distinct from integration could potentially play an important role in our experiments, and could perhaps account in part for the CA results; this mechanism is selection history. Perceptually guided choices are often influenced by the history of events recently experienced. The associated biases may be induced by past stimuli, past rewards, or past actions, for instance, or by specific combinations (Fecteau and Munoz, 2003; Awh et al., 2012; Failing and Theeuwes, 2018; Jiang and Sisk, 2019; Theeuwes, 2019; Anderson et al., 2021). Importantly, under certain experimental designs, such biases may explain effects attributed to multisensory integration (Otto and Mamassian, 2012; Shaw et al., 2020; Roberts et al., 2024).

We investigated the potential contribution of selection history effects to our results by generating history-conditioned tachometric curves for each of our task variants (Methods). For this, trials of each type (A, V or multisensory) were sorted according to their preceding trial type (again, A, V or multisensory). The main question was whether repetition of a given cue modality (e.g., V → V) reinforced performance, and conversely, whether switching from one cue modality to another (e.g., A → V) hindered performance.

In all cases, differences between history-conditioned tachometric curves were either minimal or nonexistent. The largest effect occurred in the IVA variant, where a consistent difference of approximately 12 ms was observed between rise points when the same cue type was repeated (e.g., V(A) → V(A)) versus when it switched (e.g., V → V(A)). This shift in PT was similar for the V and V(A) conditions. In the other task variants, no notable differences were found. We conclude that the contribution from history-driven effects to the current results was, for the most part, negligible.

## Discussion

In combination with urgency, the top-down control requirements of the antisaccade task produce a behavioral metric of high temporal resolution such that stimulus-driven and goal-driven contributions to performance can be dissociated. We used this unique psychophysical tool to investigate how visual and auditory cues can jointly guide saccadic choices, whether synergistically or competitively. The experiments yielded three main results.

First, the interaction between auditory and visual information evolved as the choice process unfolded. This evolution was rich and fast, reminiscent of the integration dynamics of neurons in the superior colliculus (Miller et al., 2017), and depended on the particular task configuration. But a key conclusion is that the nature of the integration process — whether performance guided by one cue is enhanced or hindered by the presence of a second cue — varies over time. An additional cue may initially enhance performance but later have no effect (Fig. 5b); it may have no effect initially but later hinder performance (Fig. 6b); or the combination may shift from a form of performance averaging to true enhancement (Fig. 3b). In general, a transition occurs between stimulus-driven and goal-driven regimes.

Contrast this with classic studies in multisensory integration in which discrimination accuracy is the main behavioral output (e.g., Ernst and Banks, 2002; Alais and Burr, 2004; Kording and Wolpert, 2004; Gori et al., 2012). Because the corresponding tasks lack urgency, not much can be inferred about the time dependence of the underlying perceptual processes, nor about the impact of fast, transient, bottom-up signals. Even in cases in which time and accuracy are considered together (Corneil et al., 2002; Drugowitsch et al., 2014), reliance on the RT obscures both the low-level contributions and the true time course of the perceptual evaluation. Non-urgent tasks sample the regime that is eventually observed after a long PT but miss the rich developments that precede it, which are highly informative of the underlying neural dynamics (Seideman et al., 2022; Zhu et al., 2024).

Second, the stimulus-driven signals derived from visual and auditory cues always played an important role; although they did not necessarily manifest in terms of ceiling accuracy (at long PTs), they always impacted the amount of PT needed to make an informed choice. To appreciate this observation it helps to realize that, for the types of sensory-guided behavior considered here, any given cue stimulus inevitably yields two signals, an exogenous one associated with its salience and an endogenous one associated with its assigned motor response (Goldstein et al., 2022, 2024; Oor et al., 2023; Zhu et al., 2024; Kattner et al., 2026). During urgent antisaccades (e.g., Fig. 2), the two signals become obvious because they point in opposite directions, one promoting a saccade to the cue and the other a saccade away. But the two components also contribute, one after the other, when they are aligned, i.e., even in the simplest case in which a participant intends to look at a salient stimulus (Oor et al., 2023; Kattner et al., 2026). The dual contribution is just more difficult to demonstrate in this case.

With this in mind, it is easier to see that in our CMC task the early stimulus-driven (exogenous) contributions were always significant and lawful. This was most evident when the cues were spatially incongruent (IVA, IAP); then the difference between unisensory performance with cue 1 and multisensory performance with cues 1 and 2 was consistent with motor plans being exogenously drawn toward cue 2 (Figs. 5b, 6b). Befitting its exogenous character, this effect occurred precisely when expected given the latency of cue 2 and regardless of the particular stimulus-response rule used. Such specific contributions were less discernible during congruent V+A trials, when both exogenous signals pointed to the one cue location and both endogenous signals pointed away. In this case, the interactions were resolved within a most compressed timescale (75,:S PT,:S 175 ms; Fig. 8b, red trace). Clearly, though, from the point of view of a visual cue that always instructed the same antisaccade, adding a congruent auditory signal yielded substantial savings in PT in comparison to an incongruent cue (∼50 ms; Fig. 8b). Such PT savings are likely a key benefit of multisensory integration more generally (Rowland et al., 2007).

In the congruent case, it is also worth noting that the change from the V to the V+A curve could not have been mimicked by a unisensory visual cue of different luminance. During urgent visual antisaccades, as stimulus luminance increases, the tachometric curve shifts to the left and the capture becomes more pronounced (Salinas et al., 2019; Goldstein et al., 2022). By contrast, the V+A tachometric curve shifted to the left while its capture became *less* pronounced (Fig. 3b red versus cyan traces). The multisensory percept was effectively less salient than the visual alone but nonetheless more effective in guiding the antisaccade. Such improvement may be qualitatively unique to cross-modal integration.

Finally, the third main finding was that multisensory performance was highly consistent with the expected allocation of attention to two cues based on the three established selection mechanisms, stimulus-driven, goal-driven, and history-driven (Corbetta and Shulman, 2002; Theeuwes, 2010, 2019; Carrasco, 2011; Awh et al., 2012; Anderson et al., 2021; Noyce et al., 2023). Endogenous selection of one cue or another was observed for identical cue-target configurations (Fig. 7b); exogenous capture was observed both when the visual cue effectively acted as a distracter (Fig. 6b) and when the auditory cue did (Fig. 5b); contextual or selection-history effects modulated the performance of visual antisaccades (Fig. 8a); and performance was less effective when attention was split between two locations than when it was focused on one (Fig. 8b).

These data speak to the relationship between attention and multisensory integration, which has been the focus of several studies (for review, see Koelewijn et al., 2010; Santangelo and Macaluso, 2012). The current consensus based on results from non-urgent tasks is that, while cross-modal attention and multisensory integration often operate independently (the traditional view), sometimes they definitely interact (a more recent view). Specifically, attention seems more likely to independently impact the integration of stimuli that are weakly associated, e.g., because they are spatially misaligned, or because they involve unrelated modality-specific features. While we did not attempt to manipulate attention and cue integration independently, our results are broadly consistent with such summary. They demonstrated both lawful low-level interactions independent of top-down goals, and strong contextual effects when cues were misaligned. Notably, whether viewed in terms of multisensory integration or crossmodal attentional allocation, results are easier to interpret when exogenous and endogenous contributions to performance are temporally dissociated.

Our experiments revealed substantial asymmetries between visual and auditory performance that are consistent with intrinsic, qualitative differences in how neural circuits process the two modalities. To be reliably produced, auditory antisaccades (A trials from CA variant) required ∼88 ms of additional PT relative to auditory prosaccades (A trials from IAP variant), a difference that is comparable to that observed with visual stimuli (Goldstein et al., 2022). However, there were two notable differences between modalities. First, overt capture was robust during visual antisaccades but minimal during auditory antisaccades (performance barely dipped below chance). And second, the asymptotic fraction correct for auditory antisaccades was considerably lower (0.81) than that obtained with visual antisaccades (0.94) or auditory prosaccades (0.95). This suggests that the cognitive operation for inverting the stimulus location and specifying the saccade target always consumed an additional amount of time (∼100 ms of PT), but in the auditory case was based on a neural signal that — in spite of the high intensity of the originating physical stimulus — was relatively weak. This is consistent with prior studies reporting strong dominance of visual information over auditory (Colavita, 1974; Spence et al., 2012; Li et al., 2017; Malevich et al., 2026).

In conclusion, the results with congruent antisaccades showed that multisensory integration is possible even when the stimulus-response rule is indirect and requires top-down control. In this case the same rule (“look away”) applied to two cues that were spatially aligned, so these may have been integrated early into a unified percept that then guided the motor selection process. In contrast, the results in the IVA/IAP experiment showed that, when spatially incongruent auditory and visual stimuli consistently signaled the same target location via different sensory-motor rules, this regularity could not be exploited by the participants to their advantage. Instead, the dynamics between incongruent multimodal cues were similar to those observed when exogenous and endogenous attention are in conflict. The distinction between these fusion and segregation modes is likely to arise in a wide variety of circumstances (Wallace et al., 2004; Magnotti et al., 2013; Shams et al., 2010; Mohl et al., 2020). Taken together, the results indicate that the capacity for integration is severely limited for stimuli that give rise to low-level spatial conflict, even when the corresponding high-level sensory-motor rules are congruent relative to the correct choice. It remains to be seen whether similar constraints apply to the integration of feature dimensions other than spatial location.

## Acknowledgments

Research was supported by the National Institutes of Health (NIH) through grants R01EY025172 and R21MH120784. We thank Denise Anderson for expert technical assistance.

## Abbreviations

A: unisensory auditory
CA: congruent antisaccades
CI: confidence interval
CMC: compelled multisensory choice
IAP: incongruent auditory prosaccades
IVA: incongruent visual antisaccades
*P_CAP_*: probability of capture
PT: processing time
RT: reaction time
V: unisen-sory visual

